# Soleus H-Reflex Up-Conditioning during Sciatic Nerve Regeneration in Rats Improves Recovery of Locomotion

**DOI:** 10.1101/2024.12.09.627519

**Authors:** Yi Chen, Yu Wang, Lu Chen, Xinxin Yang, Darren E. Gemoets, Jonathan S. Carp, Xiang Yang Chen, Jonathan R. Wolpaw

## Abstract

Operant conditioning of the spinal stretch reflex or its electrical analog, the H-reflex, induces plasticity in the brain and spinal cord that increases (up-conditioning) or decreases (down-conditioning) the reflex elicited by primary afferent input to the spinal motoneuron. In rats in which the sciatic nerve is transected and repaired, soleus (SOL) H-reflex up-conditioning during regeneration strengthens primary afferent reinnervation of SOL motoneurons and improves recovery of the SOL H-reflex. This suggests that H-reflex up-conditioning could improve functional recovery after nerve injury and repair. To explore this possibility, we examined the impact of SOL H-reflex up- or down-conditioning during sciatic regeneration on recovery of locomotor symmetry. Sprague-Dawley rats were implanted with EMG electrodes in right SOL and a stimulating cuff on right posterior tibial nerve. After control data collection, right sciatic nerve was transected and repaired. Control EMG and H-reflex data collection continued for 20 more days. The rat was then exposed for 100 days to either: continued control data collection; SOL H-reflex up-conditioning; or SOL H-reflex down-conditioning. Locomotor EMG, H-reflex, and kinematics were assessed before nerve transection and 120 days after transection. H-reflex up-conditioning improved H-reflex recovery and also restored right/left step symmetry. H-reflex down-conditioning did not worsen H-reflex recovery or right/left step asymmetry. These results suggest that H-reflex up-conditioning might enhance functional recovery after nerve injury in humans. They also confirm previous results indicating that compensatory plasticity prevents inappropriate H-reflex conditioning (i.e., down-conditioning) from further impairing function.

## INTRODUCTION

Peripheral nerve injuries are a common feature of limb trauma. With appropriate surgical care, peripheral motor and sensory axons that have been cut or crushed can regenerate and reinnervate peripheral targets (Kline and Hudson 1995; Birch 2013). However, reinnervation usually fails to reestablish the extent and specificity of connections essential for normal function: only 10% of patients regain normal motor function after traumatic transection and surgical suture of a major peripheral nerve (Frostick et al. 1998; Brushart 1998; Scholz et al. 2009).

Operant conditioning of the spinal stretch reflex or its electrical analog, the H-reflex, changes brain and spinal cord (Wolpaw and Chen, 2009; Norton and Wolpaw 2018 for review). In rats and humans with incomplete spinal cord injury, appropriate reflex conditioning improves locomotion (Chen et al., 2006b & Thompson et al., 2013). In rats in which the sciatic nerve has been transected and repaired, soleus (SOL) H-reflex up-conditioning during regeneration improves recovery of EMG activity and SOL H-reflex; this is accompanied by anatomical evidence of improvement in the extent and the specificity of primary afferent reinnervation of SOL motoneurons (Chen et al., 2010). Encouraged by these results, the present study tested the hypothesis that SOL H-reflex conditioning during sciatic regeneration also improves recovery of locomotion.

To test this hypothesis, rats were implanted with EMG electrodes in right SOL muscle and a stimulating cuff on right posterior tibial nerve. After control data collection, right sciatic nerve was transected (TX) and repaired. Data collection continued for 20 more days. The rat was then exposed for 100 days to either: SOL H-reflex up-conditioning (TU); SOL H-reflex down-conditioning (TD); or continued control data collection (TC). We assessed SOL H-reflexes and locomotor kinematics before transection (Pre TX), 20 days after transection (Post TX20), and 120 days after transection (Post TX120). The results indicate that H-reflex up-conditioning improved locomotor recovery after peripheral nerve injury and repair. Thus, they suggest that H-reflex conditioning, a noninvasive clinically practical therapy, may be able to enhance functional recovery after nerve injury in humans.

## METHODS

Subjects were 19 young male Sprague-Dawley rats with body weights of 274-455 g (mean 374(±40SD) g) at the beginning of study. All procedures were in accord with the “Guide for the Care and Use of Laboratory Animals” (National Academies Press, Washington, D.C., 2011). They had been reviewed and approved by the Institutional Animal Care and Use Committees of the Wadsworth Center and the Stratton VA Medical Center. The protocols for implantation of the nerve stimulating cuff and EMG recording electrodes, M-wave and H-reflex elicitation, chronic monitoring and conditioning of the SOL H-reflex in freely moving rats, sciatic nerve transection and repair, recording and analysis of locomotor H-reflexes and kinematics, and animal perfusion and tissue preparation have been described in detail previously (Wolpaw et al., 1993; Chen and Wolpaw, 1995, 1997, 2002, 2005, 2012; Chen et al., 2002, 2003, 2005, 2006a, 2006b, 2010, 2011, 2014a, 2014b; English, 2005; English et al., 2007; Pillai et al. 2008; Wang et al. 2006, 2009, 2012; Wolpaw and Chen, 2006; Wolpaw et al., 1993). They are summarized here.

### Implantation of Electrodes

Under general anesthesia (ketamine HCl (80 mg/kg) and xylazine (10 mg/kg), intraperitoneal, supplemented as needed) and aseptic conditions, each rat was implanted with chronic stimulating and recording electrodes in the right leg (Chen and Wolpaw, 1995, 1997, 2002; Chen et al., 2001, 2002, 2005, 2006a). To elicit the H-reflex in the right soleus muscle (SOL), a silicone rubber nerve cuff containing a pair of stainless steel multistranded fine wire electrodes was placed around the right posterior tibial (PT) nerve just above the triceps surae branches. To record right SOL EMG activity, a pair of multistranded stainless steel fine-wire electrodes (with final 0.5-cm segments exposed and separated by 0.2-0.3 cm) were placed in the right SOL muscle. The Teflon-coated wires from the muscle and the nerve cuff passed subcutaneously to a connector secured to the skull with stainless steel screws and dental cement. In addition, small (2-mm) dots were tattooed bilaterally on the skin overlying the lateral aspects of the knees, hips (i.e., trochanter major), and iliac crests at the 5th lumbar vertebra. These marks were used to guide consistent placement of reflective markers during video recording of treadmill locomotor sessions (i.e., Vicon System, see below and Figure 1c); they facilitated repeated kinematic analyses over the prolonged period of data collection. After surgery, the rat was placed under a heating lamp and given an analgesic (Demerol, 0.2 mg, i.m.). Once awake, it received a second dose of analgesic and was returned to its home cage. A high-calorie dietary supplement was given until body weight regained its pre-surgery level. Each rat also received a piece of apple (∼10 g) every day. All the rats gained weight and remained healthy and active throughout study.

**Figure 1.**
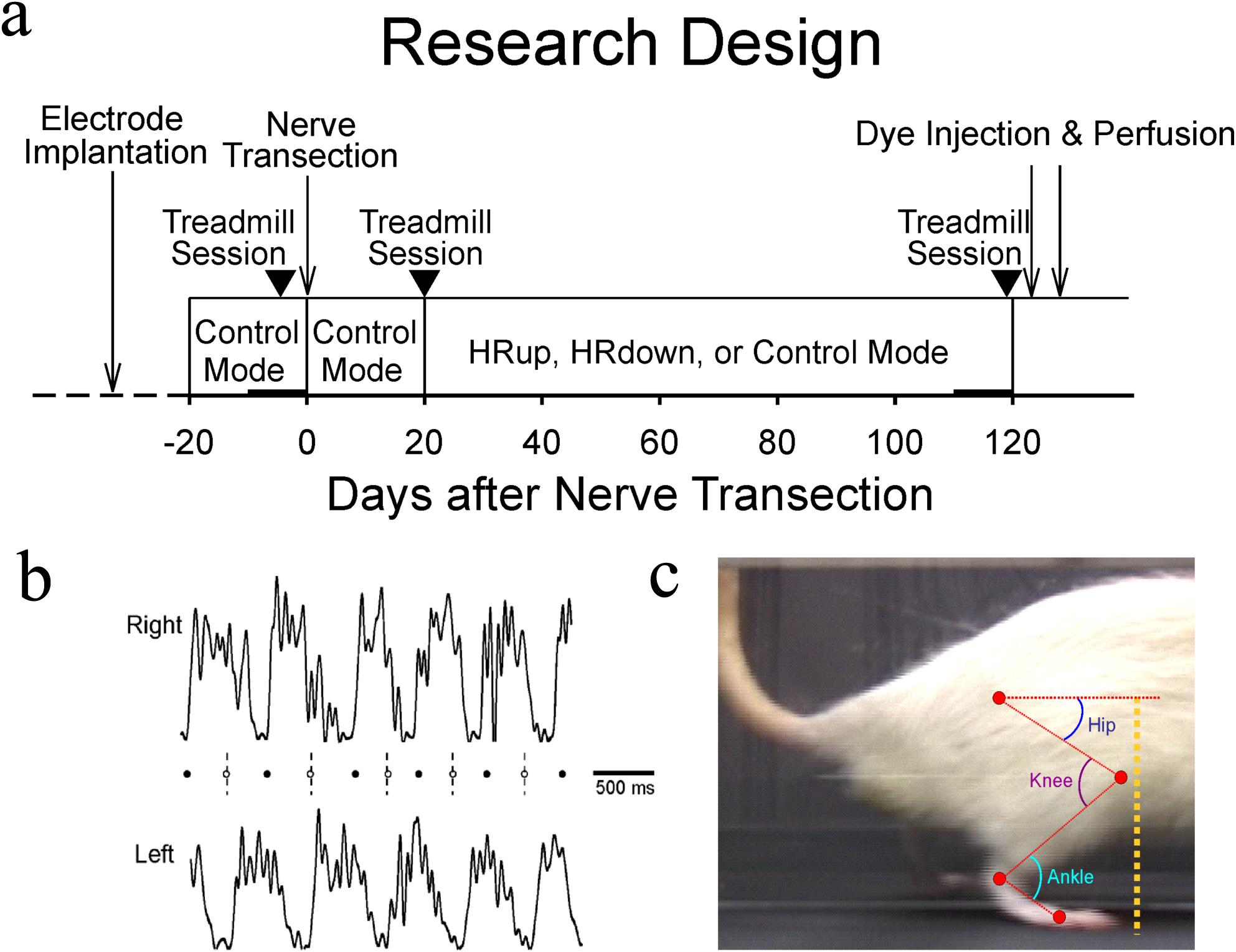
Experimental design and methodology. **Panel 1a**: Sprague-Dawley rats were implanted with EMG electrodes in the right soleus muscle and a stimulation cuff on the right posterior tibial nerve. After 20 days of control H-reflex data collection (Control Mode), the right sciatic nerve was transected and repaired. After 20 more days of Control-Mode H-reflex data collection, each rat was then exposed for 100 days to either: continued Control Mode data collection (TC rats); soleus H-reflex up-conditioning (HRup) (TU rats); or soleus H-reflex down-conditioning (HRdown) (TD rats). Locomotor EMG, H-reflex, and kinematics were assessed before transection, and 120 days after transection. **Panel 1b**: Assessing locomotor function during treadmill walking in rats. Rectified right (upper trace) and left (lower trace) soleus bursts during walking from a normal rat. In normal rats, the time from the right foot contact (filled circles) to the left foot contact (open circles) equals that from the left foot contact to the right foot contact, i.e., the step cycle is symmetrical, as indicated by the dashed vertical lines being midway between right foot contacts. **Panel 1c**: Illustration of measurement of ankle, knee, and hip angles, and hip height, i.e., vertical distance from the hip to the ground (dotted yellow line).

### Transection and repair of the sciatic nerve

After 20 days of control data collection (which started at least 20 days after the implantation surgery) (Figure 1a), each rat underwent a second surgery in which the right sciatic nerve was transected and repaired as described in English et al. (2007) and Chen et al. (2010). Briefly, under general anesthesia and aseptic conditions, the sciatic nerve was exposed proximal to the branching of the sural nerve and cut completely with sharp scissors. The distal stump was immediately aligned with the proximal stump using surface landmarks such as blood vessels and connective tissues as a guide. The two stumps were then glued together with fibrin glue (English, 2005; English et al., 2007; Chen et al. 2010). Post-surgical care was as described above.

### Data collection and H-reflex conditioning

Figure 1a shows the experimental design. To study the impact of SOL H-reflex conditioning during sciatic regeneration on H-reflex function and locomotion, electrophysiological data collection began at least 20 days after the implantation surgery and continued 24 hrs/day, 7 days/week for 140 days as regeneration was occurring (English et al. 2007; Chen et al. 2010). During this period, each rat lived in a standard rat cage with a 40-cm flexible cable attached to the skull plug. The cable, which allowed the animal to move freely about the cage, conveyed the wires from the electrodes to a commutator above it, and from there to EMG amplifiers (gain 1000, bandwidth 100-1000 Hz) and a stimulus isolation unit. The rats had free access to water and food, except that during H-reflex conditioning they received food mainly by performing the task described below. Animal well-being was carefully checked several times each day, and body weight was measured weekly. Laboratory lights were dimmed from 2100 to 0600 each day (Chen and Wolpaw 1995, 1996, 1997, 2002, 2005, 2012; Chen et al. 2001, 2003, 2005, 2006a, 2006b, 2010, 2011).

Stimulus delivery and data collection were under the control of a computer system, which monitored ongoing SOL EMG activity continuously 24 hr/day, 7 days/week, for the entire period of data collection. The EMG signal was sampled at 5,000 Hz and bandpass filtered at 100-1,000 Hz. Whenever the absolute value (equivalent to the full wave rectified value) of SOL background (i.e., ongoing) EMG activity remained within defined ranges for a randomly varying 2.3-2.7 s period, the computer initiated a trial. In each trial, the computer stored the most recent 50 ms of EMG activity from the muscle (i.e., the background EMG interval), delivered a monophasic stimulus pulse to the nerve cuff, and stored the EMG activity for the following 100 ms. The amplitude and duration of stimulus pulses were initially set to produce a maximum H-reflex (as well as an M-wave that was typically just above threshold). The pulse duration remained fixed (usually 0.5 ms), while the pulse amplitude was adjusted by the computer after each trial to maintain SOL M-wave size. The M-wave size was defined as the average absolute value of EMG activity within the M-wave interval, usually 2.0-4.5 ms after PT nerve stimulation. This approach ensured that the effective strength of the nerve stimulus was stable throughout the experiment, despite any changes that occurred in nerve cuff electrode impedances or in other factors (Wolpaw, 1987; Chen and Wolpaw, 1995).

During the period after the nerve transection and before the M-wave had fully recovered, the stimulus pulse was permitted to rise to 2.5 times its average pre-transection amplitude (i.e., average amplitude in the final 10 days before the nerve transection), or the amplitude required to achieve the target M-wave size, whichever was lower. Also, during this period in which the M-wave gradually returned and its latency shortened, the M-wave interval was repeatedly adjusted appropriately. The H-reflex interval was similarly adjusted over this period. Typically, both responses regained their initial pre-transection latencies, and the M-wave regained its pre-transection size by 80-100 days after transection. This was well before the end of data collection, when final H-reflex size was determined, and final locomotor performance was assessed (described in the next subsection). H-reflex size was defined as average absolute amplitude of EMG activity in the H-reflex interval (typically 6-10 ms after PT nerve stimulation) minus average background EMG amplitude (Chen and Wolpaw 1995, 1996, 1997, 2002, 2005, 2012; Chen et al. 2001, 2003, 2005, 2006a, 2006b, 2010, 2011).

Under the control mode of the H-reflex conditioning protocol, the computer simply digitized and stored the absolute value of SOL EMG activity for 100 ms following the stimulus as described above. Under the up-conditioning or down-conditioning modes, it gave a reward (i.e., a food pellet) 200 ms after PT nerve stimulation if SOL EMG activity in the H-reflex interval was above or below a criterion value, respectively. The criterion value was set initially and adjusted as needed each day, so that the rat received an adequate amount of food and body weight was maintained (e.g., about 700-900 reward pellets per day for a 400-gram rat). As shown in Figure 1a, each rat was first studied under the control mode for 20 days. The data for the final 10 days of this period were averaged to provide the initial values of pre-transection (Pre TX) H-reflex size. The rats then underwent sciatic nerve transection and repair. Following this surgery, data collection continued for 120 more days to assess the impact of H-reflex conditioning during sciatic regeneration. During this 120-day period, 7 transected/control (TC) rats simply continued under the control mode (i.e., prolonged control for a total of 120 days); 6 transected/up-conditioned (TU) rats continued under the control mode for 20 days and were then exposed to the up-conditioning mode for the remaining 100 days; and 6 transected/down-conditioned (TD) rats continued under the control mode for 20 days and were then exposed to the down-conditioning mode for the remaining 100 days.

To determine the impact of H-reflex conditioning during sciatic regeneration on H-reflex size, average daily H-reflex size for the final 10 days Post TX120 (i.e., Days 111-120 following the nerve transection) of continued control mode, up-conditioning, or down-conditioning (TD_120_ rats) (thick x-axis segment on the right in Figure 1a), was expressed in percent of the average daily H-reflex size for the final 10 control-mode days before the nerve transection (Pre TX; thick x-axis segment on the left in Figure 1a). These H-reflexes are referred to as the ‘protocol H-reflexes’ because they were elicited and evaluated during the 24/7 H-reflex conditioning protocol. They are distinguished from the ‘locomotor H-reflexes,’ which were elicited during treadmill walking sessions as described below.

Finally, we assessed the activity level of each rat over the period of regeneration as the average number of protocol H-reflexes per day multiplied by the average level of soleus background EMG prior to each trial.

### Locomotor Data Collection

Before the initial surgery (i.e., electrode implantation, Figure 1a), each rat walked quadrupedally on a motor-driven treadmill at 9-12 m/min for three or four 15-25 min training sessions (Chen et al., 2005, 2006b, 2011, 2014a, 2014b, 2017). Locomotor data were then collected from each rat in the three treadmill sessions, as shown in Figure 1a: before sciatic nerve transection, 20 days after transection, and 120 days after transection. In each rat, treadmill speed remained the same for the three sessions.

Prior to each locomotor session, the hindlimbs were shaved and 3-mm reflective adhesive markers were placed (aided by the tattooed dots (see above)) on the lateral aspects of the fifth metatarsophalangeal joint, the ankle joint (i.e., lateral malleolus), the knee joint, the hip joint (i.e., trochanter major), the iliac crest at the 5th lumbar vertebra, and the midpoint between the ankle and knee joints of each leg. These markers enable later analysis of locomotor kinematics.

During locomotion, SOL EMG activity was continuously recorded, bandpass filtered (100-1000 Hz), digitized (5,000 Hz), and stored. In addition, locomotor kinematics were recorded bilaterally with a 3-D video data collection and analysis system (100 frames/sec) (Vicon Motion Systems). In each treadmill session, data were collected under two conditions. One was undisturbed locomotion. In the other, the SOL H-reflex was elicited by stimulating the PT nerve just after the middle of the stance phase (i.e., the ‘locomotor H-reflex’) as described in Chen et al. (2005, 2006a, 2014a, 2014b, 2017). About 5 min (i.e., 400∼500 step cycles) of data were collected under each condition.

### Analysis of Locomotor Data

As done for the size of the H-reflex elicited in the conditioning protocol (i.e., the ‘protocol H-reflex’), the size of the H-reflex elicited during locomotion (i.e., the locomotor H-reflex) was calculated as average absolute value of EMG activity in the H-reflex interval minus average absolute value of background EMG activity at the time of stimulation and was expressed in units of average absolute value of background EMG activity (See Chen et al. (2005) for full description of this measure). The effects of H-reflex conditioning on the locomotor H-reflex were assessed by comparing the reflexes before and after up-conditioning, down-conditioning, or continued control-mode exposure.

The concurrent 3-D locomotor kinematic data were analyzed with the Vicon Motus software (Vicon Motion Systems) to assess two measures that reflect the right/left symmetry of the step cycle. The first measure was step symmetry, defined as the average time from right foot contact (RFC) to left foot contact (LFC) divided by the average time from LFC to RFC. It was expressed in percent of the average time from LFC to RFC (e.g., 100% indicates that the step cycle is perfectly symmetrical). The second measure was hip-height symmetry, defined as the average right hip height (e.g., Figure 1b) divided by the average left hip height, and expressed in percent of the average of right and left hip height (e.g., 100% indicates that the hip heights are symmetrical).

### Perfusion, Postmortem Examination, and Lesion Verification

At the end of the study, the rat received an overdose of sodium pentobarbital (i.p.) and was perfused through the heart with saline followed by 4% paraformaldehyde in 0.1 M phosphate buffer (pH 7.3). The EMG electrodes, nerve cuff, and PT nerve were examined, and the SOL muscles of both sides were removed and weighed.

## RESULTS

Animals remained healthy and active throughout the study. Body weight increased from 374(±40SD) g at the time of implantation surgery, to 516(±61) g at the time of nerve transection, and to 697(±92) g at the time of perfusion. The control (TC), up-conditioned (TU), and down-conditioned (TD) rat groups did not differ significantly in their weight gains over the period of study (p=0.89 by one-way ANOVA). Postmortem examination of the nerve cuffs revealed the expected connective tissue investment of the fine-wire electrodes and good preservation of the PT nerve within this connective tissue sheath. The right SOL muscle (i.e., the sciatic-transected hindlimb) weighed 0.20 (±0.04 SD) g, significantly less than the left SOL muscle (i.e., the intact hindlimb), which averaged 0.26 (±0.03) g (p<0.001 by paired t-test). The weights of the right and left SOL muscles did not differ significantly across the TC, TU, and TD rat groups (p=0.23 and p=0.69 by One-way ANOVA for right and left SOL, respectively).

### Level of Activity and Number of Rewards

As described in the Methods section, whenever the background EMG requirement was satisfied, nerve cuff stimulation elicited the M-wave and H-reflex. Thus, the overall activity level of the rat was reflected by the product of the average number of elicitations (trials) per day and the average background EMG level (absolute value in µV) immediately before a trial. Table 1 shows the average value of this measure before transection and for the 120 days after transection for each of the three rat groups. It also shows for the TU and TD rat groups, the average number of rewards (i.e., pellets)/day for the conditioning period. As indicated there, the activity levels of TC, TU, and TD rats did not differ significantly before transection, but did differ after transection (p<0.001); activity level was higher in TU rats than in TD or TC rats (p=0.004 and p<0.001, respectively). TU and TD rats did not differ in the number of rewards (i.e., pellets) they received per day.

**Table 1:** Activity Level (trials/day x Background EMG level (uV)) before and after nerve transection for each rat group, and Rewards/Day for the H-reflex conditioning groups.

| Rat Groups | Activity Level (trials/day x EMG level (uV)) |  | Rewards/Day |
| --- | --- | --- | --- |
|  | Before transection | Days 1-120 after transection |  |
| Up Group (N=6) | 399453 | 500486 | 1433 |
| Down Group (N=6) | 490382 | 261572 | 1530 |
| Control Group (N=7) | 499351 | 169634 | - |

Because these intergroup differences could conceivably contribute to the functional differences among the groups that are described below, we also performed an Analysis of Covariance (ANCOVA) to determine whether the functional differences could be ascribed to the differences in average activity level during the 120-day post-transection period (i.e., Table 1). Similar results were obtained using an ANCOVA. Thus, the final differences in function could not be ascribed to differences in average activity level.

### Background EMG activity

EMG activity in the right SOL fell immediately after transection. Thus, to permit H-reflex trials, the minimum background EMG requirement was lowered. It was gradually raised as regeneration occurred and SOL EMG activity increased. Figure 2a shows, for TU, TD, and TC rat groups, average SOL background activity for the final 10 days of data collection (111-120 days after transection in Figure 1a) in percent of its pre-transection value. In TU rats, final post-transection background activity did not differ from the pre-transection value (p=0.16 by paired t-test). In TC and TD rats, final post-transection background activity was significantly lower than its pre-transection value (p<0.001 and p=0.009, respectively). In sum, background EMG returned to near its background level only in up-conditioned (TU) rats.

**Figure 2.**
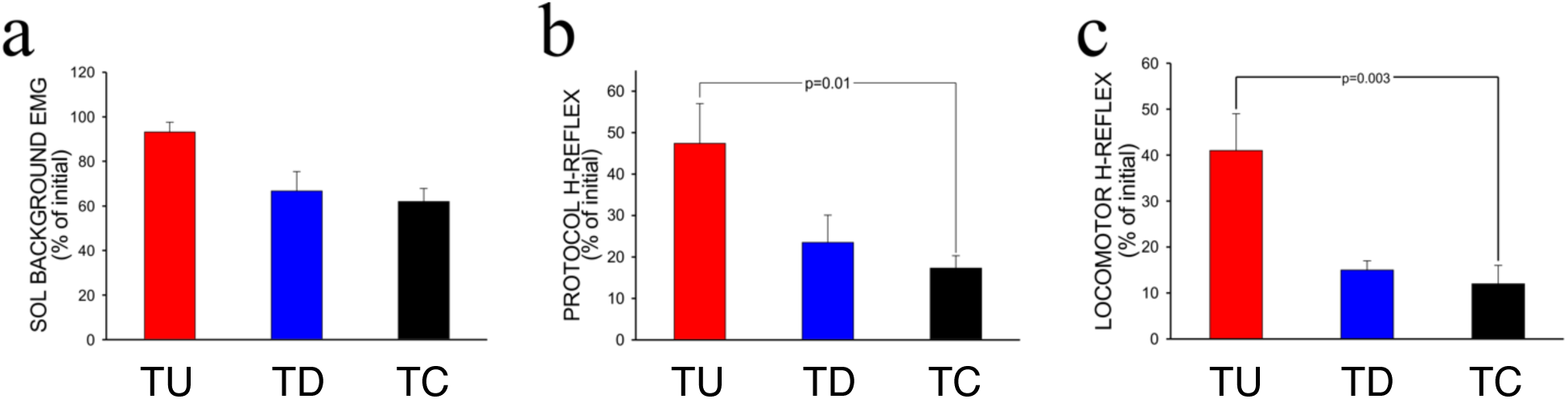
Effect of H-reflex up-conditioning (TU rats, n=6), H-reflex down-conditioning (TD rats, n=6), and continued Control-Mode (TC rats, n=7) on the average soleus background EMG (panel a), protocol H-reflex (panel b), and locomotor H-reflex (panel c) for the final 10 days of data collection (111-120 days after transection (Figure 1a)) as a percent of its pre-transection value. Differences were assessed by ANOVA with post-hoc Tukey’s HSD test (see text for details of statistical analyses). **Panel a**: Background EMG recovered to near its background level in TU rats, but not in TC or TD rats. **Panel b**: In all three groups, the final protocol H-reflexes were significantly lower than their pre-transection values; however, a greater degree of recovery of the protocol H-reflex was seen in TU rats than in TC rats. **Panel c**: As seen with the protocol H-reflex, the final locomotor H-reflex exhibited a greater degree of recovery in TU rats than in TC rats.

### Protocol and Locomotor H-reflexes

The right SOL H-reflex elicited whenever the background EMG requirement was met (i.e., the protocol H-reflex) disappeared immediately after sciatic nerve transection. It reemerged several weeks later and grew gradually over the next several months. Figure 2b shows for each rat group average SOL H-reflex size for the final 10 days of data collection in percent of its pre-transection value. In all three groups, the final value was significantly lower than the pre-transection value. At the same time, the three groups differed significantly (p=0.013 by ANOVA). The final value was significantly greater in TU than in TC (p=0.01; Tukey HSD method) rat groups but did not differ significantly between TD and TC rats (p=0.78). The average final H-reflex size in TU rats was more than twice that in TD or TC rats.

Finally, in each rat group the effect of transection on the final locomotor H-reflex (Figure 2c) was similar to its effect on the final protocol H-reflex (Figure 2b). In all the rats, the final protocol and locomotor H-reflexes were significantly correlated (r=0.73). The three groups differed significantly (p=0.002 by ANOVA); and the final locomotor H-reflex in the TU rats was significantly larger than that in the TC rats (p=0.003; Tukey HSD test). TC and TD rats did not differ significantly (p=0.81).

### Locomotor Kinematics

Figure 3 summarizes the impact of transection on locomotor kinematics. Average step length was minimally affected. In contrast, transection greatly affected step symmetry—the right step became substantially shorter than the left. By the end of data collection (i.e., 120 days after transection), step symmetry had been almost fully restored in TU rats (p = 0.61 vs. pre-transection value). It remained asymmetrical in TD and TC rats (p=0.02 in both), although it had improved considerably in TD rats (p=0.03 vs. 20 days after transection). In sum, H-reflex up-conditioning restored step symmetry, down-conditioning improved it somewhat, and the control mode had no detectable effect.

**Figure 3.**
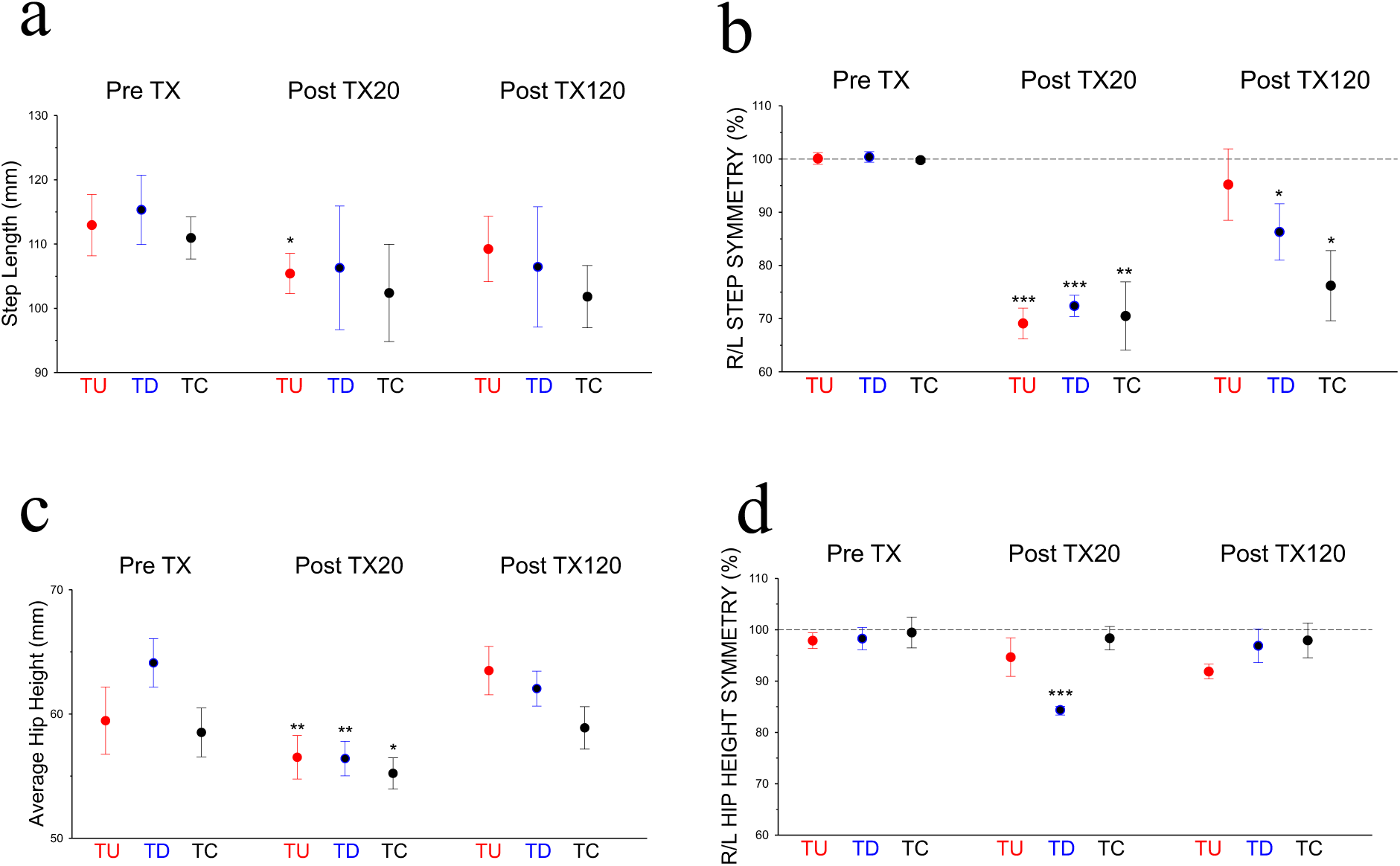
Hindlimb kinematic properties before (Pre TX), 20 days (Post TX20), and 120 days (Post TX120) after sciatic nerve transection and repair in rats exposed to H-reflex up-conditioning (TU rats, n=6), H-reflex down-conditioning (TD rats, n=6), and continued Control-Mode (TC rats, n=7) for 100 days immediately following the Post TX20 recording session. Differences between Pre TX, Post TX20, and Post TX120 values were assessed by ANOVA with post-hoc Tukey’s HSD test within each of the TU, TD, and TC treatment groups (*p<0.05, **p<0.01, ***p<0.001 for Pre TX vs. Post TX20 or Post TX120; see text for details of statistical analyses). **Panel a**: Locomotor step length was not significantly affected by H-reflex conditioning treatments at either the Post TX20 or TX120 time points. **Panel b**: Nerve transection (Post TX20) resulted in a significant decrease in step length on the injured side, resulting in a marked asymmetry in right-left step length. TU rats recovered symmetric locomotion (not significantly different from the Pre Tx condition). TD and TC rats exhibited only partial recover of locomotor step symmetry (their step symmetry ratios were still significantly different from the Pre Tx condition. **Panel c**: The average of left and right hip height (panel c) decreased significantly after nerve transection (Post TX20); after 100 days of TU, TD, or TC (Post TX120), hip height recovered (no significant difference between the Pre TX and Post TX120 time points). **Panel d**: The ratio of right to left hip height (R/L hip height symmetry) decreased after nerve transection in the TD rat group, but not in the other (TU, TC) groups prior to 100-day exposure to HR operant conditioning or control mode stimulation. Hip height symmetry in all three treatment groups after TU, TD, or TC (Post TX120) were not significantly different from that seen prior to nerve transection (Pre TX).

Average hip height (i.e., average of right and left) did not differ among the three groups before transection (Pre TX), 20 days after transection (Post TX20), or 120 days (Post TX120) after transection (p=0.20, 0.78, and 0.17, respectively, by ANOVA; Figure 3c). At the same time, average hip height in all three groups was significantly decreased at Post TX20 (p=0.003, 0.003, and 0.02, respectively by paired t-test) and had recovered by Post TX120 (p=0.18, 0.26 and 0.72, respectively).

Before transection (Pre TX), right and left hip heights did not differ significantly in any of the three groups (p=0.24, 0.39, and 0.77 in TU, TD, and TC rats, respectively, by paired t-test). The slightly (but not significantly) lower right hip height evident in Figure 3d might reflect the presence of the EMG electrodes and nerve cuff in the right leg (hip height as % of pre-transection control was 97.9(±1.5)%, 98.3(±2.2)%, and 99.5(±3.0)% (p=0.24, 0.39, and 0.77) in TU, TD, and TC rats, respectively, by paired t-test). Twenty days after transection (i.e., before treatment of the three groups diverged (Figure 1a)), right hip height was significantly lower in the TD group (p<0.001), but not in the TU or TC group (p=0.23 and 0.44, respectively. By 120 days after transection, right and left hip heights did not differ significantly in any group (p=0.06, 0.20, and 0.47, for TU, TD, and TC rats, respectively).

## DISCUSSION

In animals and humans, transected peripheral nerves that are surgically repaired so that the proximal and distal stumps are properly opposed to each other will regenerate, and sensorimotor function will recover to some degree. However, functional recovery is usually incomplete: nine of ten patients who experience a major peripheral nerve transection and undergo surgical suture are left with functional impairment, even after regeneration has occurred (Frostick et al. 1998; Brushart 1998; Scholz et al. 2009). The persistent functional deficits are associated with and are believed to be caused by imprecise regeneration; axons often fail to reach their appropriate targets and may instead reach inappropriate targets (Sunderland, 1978; Brushart and Mesulam, 1980; Brushart et al., 1983; Fu and Gordon, 1995; Edgerton et al., 1997; English, 2005). Thus, new methods for improving the specificity of regeneration could enhance functional recovery.

Operant conditioning protocols can target plasticity to specific spinal reflex pathways, such as the largely monosynaptic pathway of the H-reflex, the electrical analog of the spinal stretch reflex (Wolpaw, 2006b; Wolpaw and Chen, 2009). They can thereby affect motor skills, such as locomotion, that use these pathways. Our previous studies examined the impact of operant up-conditioning of the soleus H-reflex during regeneration of a transected sciatic nerve (Chen et al., 2010). We found that up-conditioning, which strengthened the spinal pathway of the H-reflex, accelerated recovery of SOL EMG activity, and increased recovery of the SOL H-reflex. Furthermore, these improvements were associated with greater numbers of primary afferent excitatory (i.e., glutaminergic) terminals on SOL motoneurons.

Encouraged by these results, and by the beneficial impact of H-reflex conditioning in humans and animals with incomplete spinal cord injury (Thompson and Wolpaw 2014, 2015; Chen et al., 2006b), the present study asked whether SOL H-reflex conditioning during sciatic nerve regeneration in rats could enhance recovery of locomotion. We studied three groups of rats: in the TU group, the SOL H-reflex was up-conditioned from Days 20-120 after transection); in the TD group, it was down-conditioned; in the TC (i.e., Control) group, it was simply measured. We studied the impact on recovery of background EMG activity, the SOL H-reflex size, and locomotor function. The results were clear and encouraging.

### Effects on SOL background activity and H-reflex

We have previously reported that up-conditioning of SOL H-reflex promoted recovery of background EMG activity, M response, and H-reflex (English et al., 2007, Chen et al., 2010).

The present study confirms these results. H-reflex up-conditioning (TU) increased recovery of background EMG and H-reflex size over that found in the Control (TC) group (Figure 2). The impact on the H-reflex was not limited to the conditioning protocol; H-reflex up-conditioning also increased the final size of the H-reflex elicited during treadmill locomotion. The lack of similar improvements in the down-conditioned (TD) rats indicates that improvement was not a nonspecific effect of conditioning; it was specific to the reward criterion.

These results cannot be explained by the known effects of electrical stimulation on axonal regeneration (Nix and Hopf, 1983; Al Majed et al., 2000; Brushart et al., 2005; Gordon et al., 2008). The stimuli that enhanced regeneration in these earlier studies were delivered to the proximal stump of the cut nerve at relatively high rates (e.g., 20 Hz) and intensities. In contrast, the stimuli of the present study were delivered to the distal stump as single pulses at a rate <0.3 Hz and at low intensity (usually just above M-wave threshold). Furthermore, and most important, the results cannot be explained by the known effects of electrical stimulation because they were specific to up-conditioning, they did not occur in down-conditioned (TD) or in unconditioned (TC) rats.

At the same time, it is interesting that down-conditioning did not worsen the results. Background EMG and H-reflex recovery were somewhat better (though not significantly different) in the TD group than in the TC group. This finding is consistent with previous results in rats with incomplete spinal cord injuries: while appropriate H-reflex conditioning improves locomotion, inappropriate H-reflex conditioning does not further impair it (Chen et al., 2014a). It appears that compensatory plasticity occurs that prevents H-reflex decrease from further impairing locomotion (Chen et al., 2014a for discussion). It is likely that adjustments by other muscles, and possibly additional adaptive activity-dependent plasticity, compensate for the change in the soleus locomotor burst to preserve normal locomotor symmetry (Chen et al., 2005).

### Effects on locomotion

Earlier studies found that appropriate H-reflex conditioning could restore step symmetry in rats and humans with incomplete spinal cord injuries (Chen et al. 2006b; Thompson et al., 2013; Thompson and Wolpaw 2014, 2015; Chen et al., 2014a). In this study, we found that unilateral transection and repair of the sciatic nerve produced asymmetric stepping in all the three rat groups (Fig 3b). The reduction in the step symmetry was comparable in the TC, TU, and TD groups. Up-conditioning of SOL H-reflex during nerve regeneration largely restored step symmetry. By 120 days after transection, the hindlimb step lengths of the TU rats were symmetrical. In contrast, step-length asymmetry persisted in the TD and TC groups up to 120 days post-transection.

Immediately after nerve transection and repair, the average hip heights in all the three groups decreased significantly (Figure 3c) and they all increased to a level that was not statistically significant from their pre-transection levels. Measurements of hip height over time are likely complicated by ongoing growth of the rat, which may have obscured differences in hip height due to the different treatments.

### Conclusions

During the period of sciatic nerve regeneration after a transection injury, SOL H-reflex up-conditioning in rats significantly increase recovery of both the SOL protocol H-reflex and the locomotor H-reflex while down-conditioning did not make recovery of them worse. Both up- and down-conditioning appeared to improve locomotor step symmetry. On the other hand, H-reflex conditioning did not have clear effects on locomotor EMG bursts or the symmetry of the right/left hip height. While the mechanisms responsible for these functional and anatomical effects are yet to be further explored, the results of this study suggest that operant conditioning of spinal reflexes could be a useful tool for modifying the outputs of spinal circuits to improve functional recovery after peripheral nerve injuries.

## Acknowledgments

We thank Drs. Sebastian Rueda Parra and Elizabeth Winter Wolpaw for valuable comments on the manuscript. This work was supported by grants from the VA (P01 HD32 (JRW)) and from NIH (HD36020 (XYC), NS22189 (JRW), NS061823 (XYC&JRW), HD032571 (Dr. Arthur W. English), NIBIB / 1P41EB018783 (JRW)) and by the Stratton VA Medical Center.

